# Rapid radiation of Southern Ocean shags in response to receding sea ice

**DOI:** 10.1101/2021.08.18.456742

**Authors:** Nicolas J. Rawlence, Alexander T. Salis, Hamish G. Spencer, Jonathan M. Waters, Lachie Scarsbrook, Richard A. Phillips, Luciano Calderón, Timothée R. Cook, Charles-André Bost, Ludovic Dutoit, Tania M. King, Juan F. Masello, Lisa J. Nupen, Petra Quillfeldt, Norman Ratcliffe, Peter G. Ryan, Charlotte E. Till, Martyn Kennedy

**Affiliations:** Department of Zoology, University of Otago, Dunedin, New Zealand; Australian Centre for Ancient DNA, University of Adelaide, South Australia, Australia; British Antarctic Survey, Natural Environment Research Council, United Kingdom; Instituto de Biología Agrícola de Mendoza (IBAM, CONICET-UNCuyo), Argentina; FitzPatrick Institute of African Ornithology, Department of Biological Sciences, University of Cape Town, South Africa; CEBC-CNRS, UMR 7372, 405 Route de Prissé la Charrière, 79360 Villiers en Bois, France; Justus Liebig University, Giessen, Germany; Organisation for Tropical Studies, Skukuza, South Africa; School of Human Evolution and Social Change, Arizona State University, Arizona, USA

**Keywords:** Biogeography, climate change, cormorant, *Leucocarbo*, speciation, Southern Ocean, sub-Antarctic

## Abstract

**Aim:** Understanding how wild populations respond to climatic shifts is a fundamental goal of biological research in a fast-changing world. The Southern Ocean represents a fascinating system for assessing large-scale climate-driven biological change, as it contains extremely isolated island groups within a predominantly westerly, circumpolar wind and current system. The blue-eyed shags (*Leucocarbo* spp.) represent a paradoxical Southern Ocean seabird radiation; a circumpolar distribution implies strong dispersal capacity yet their speciose nature suggests local adaptation and isolation. Here we use genetic tools in an attempt to resolve this paradox.

**Location:** Southern Ocean.

**Taxa:** 17 species and subspecies of blue-eyed shags (*Leucocarbo* spp.) across the geographical distribution of the genus.

**Methods:** Here we use mitochondrial and nuclear sequence data to conduct the first global genetic analysis of this group using a temporal phylogenetic framework to test for rapid speciation.

**Results:** Our analysis reveals remarkably shallow evolutionary histories among island-endemic lineages, consistent with a recent high-latitude circumpolar radiation. This rapid sub-Antarctic expansion contrasts with significantly deeper lineages detected in more temperate regions such as South America and New Zealand that may have acted as glacial refugia. The dynamic history of high-latitude expansions is further supported by ancestral demographic and biogeographic reconstructions.

**Main conclusions:** The circumpolar distribution of blue-eyed shags, and their highly dynamic evolutionary history, potentially make *Leucocarbo* a strong sentinel of past and ongoing Southern Ocean ecosystem change given their sensitivity to climatic impacts.

## 1. INTRODUCTION

Understanding how wild populations respond to climatic shifts represents a key goal of biological research in a fast-changing world (Parmesan & Yohe, 2003; Chen et al., 2011). Climate change can underpin major shifts in the distribution and diversity of biological assemblages, with glacial-interglacial cycles thought to have driven dramatic changes in species distributions across many regions of the globe (Davis & Shaw, 2001). The biogeographic responses of Northern Hemisphere biota to Quaternary climate shifts, and associated ice sheet dynamics, are relatively well studied (Hewitt, 2000; Maggs et al., 2008). By contrast, climate-driven shifts involving Southern Hemisphere taxa, especially those inhabiting the Southern Ocean, remain less understood (Fraser et al., 2009; Fraser et al., 2010; Fraser et al., 2012; Cole et al., 2019).

The Southern Ocean represents a fascinating system for assessing large-scale climate-driven biological change, as it contains multiple groups of extremely isolated oceanic islands within the predominantly westerly wind and current system encircling Antarctica (Fraser et al., 2012). Understanding the evolutionary dynamics of these unique island ecosystems, and of their distinctive endemic taxa, is particularly important in the context of the region’s dynamic climatic history. Specifically, geological and biological evidence together indicate that, during recent glacial maxima, several of these outlying island groups (e.g., South Georgia, Kerguelen, Heard, Crozet, Macquarie [itself only ~700,000 years old]) were encompassed by winter sea ice for thousands of years (Fig. 1) (Gersonde et al., 2005; Fraser et al., 2009; Trucchi et al., 2014) and in some cases potentially covered in ice and snow (Hodgson et al., 2014). Indeed, Heard Island was permanently covered in ice during the Last Glacial Maximum (Ehlers et al., 2011).

**Figure 1.**
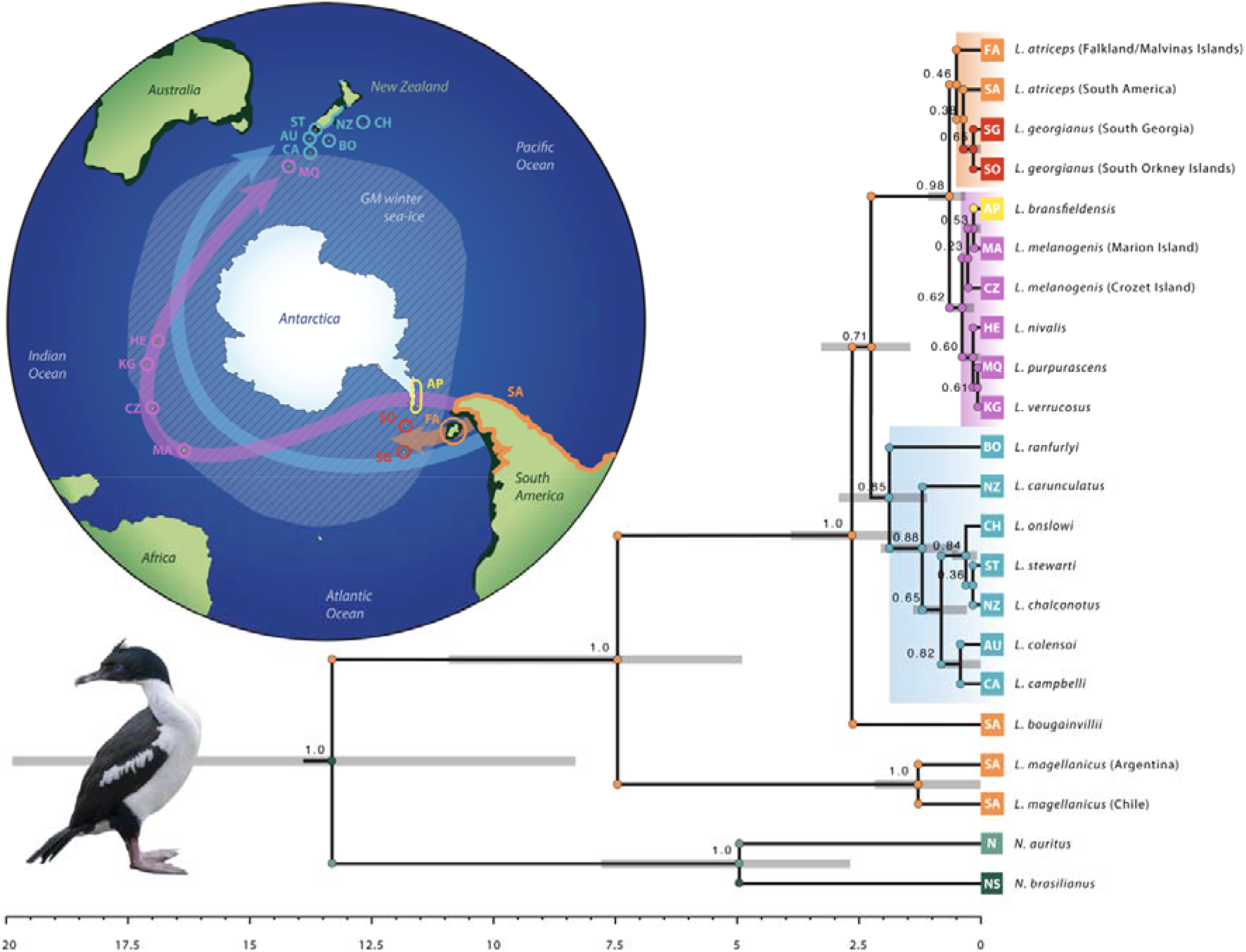
Evolutionary history of blue-eyed shags (*Leucocarbo* spp.) in the Southern Ocean. The map depicts inferred postglacial colonization routes (arrows) supported by the dominant westerly winds and eastward flow of the Antarctic Circumpolar Current. The extent of winter sea ice (cross-hatched pattern) and land area (dark green) during the Pleistocene Last Glacial Maximum 29-19 Kya is indicated. Rapid circumpolar expansion and founder speciation hypotheses are supported by temporal phylogenetic and ancestral biogeographic reconstructions. The time-calibrated species tree is derived from 8.2 kilobases of DNA sequence data (five mitochondrial and five nuclear genes) with *Nannopterum* as the outgroup. Node bars on the phylogeny are 95% HPD of divergence times as indicated on the scale bar (millions of years before present). Node values are Bayesian posterior probability support. Node circles are the ancestral state reconstruction of the geographic distribution based on the DEC+J model. Colours and abbreviations are as follows: orange: South America; purple: high-latitude sub-Antarctic islands; yellow: Antarctic Peninsula; blue: New Zealand region; olive: North America; dark green: South America; FA: Falkland/Malvinas Islands; SA: South America; SG: South Georgia; SO: South Orkney Islands: AP: Antarctic Peninsula; MA: Marion Island; CZ: Crozet Island; HE: Heard Island; MQ: Macquarie Island; KG: Kerguelen Island; CA: Campbell Island; AU: Auckland Island; NZ: mainland New Zealand; ST: Stewart Island; CH: Chatham Islands; BO: Bounty Islands; N: North America; NS: North and South America. Clade and arrow colours: orange: South America, South Georgia, South Orkney Islands; purple: Antarctic Peninsula, high-latitude sub-Antarctic islands; blue: New Zealand region.

The Pleistocene Ice Age (2.58 – 0.01 Mya) was characterized by repeated glacial-interglacial cycles (Ehlers et al., 2011). Major deglaciation events can provide significant ecological opportunities for surviving species (Fraser et al., 2018). Indeed, recent genetic data suggest that some highly-dispersive Southern Ocean species or communities have, at different timescales, responded rapidly to colonize newly vacant habitats arising from such climate-driven ecosystem change (de Bruyn et al., 2009; Fraser et al., 2009; Fraser et al., 2010; Nikula et al., 2010; Fraser et al., 2011; Fraser et al., 2012; Fraser et al., 2018; Gonzalez-Wevar et al., 2017; Gonzalez-Wevar et al., 2018; Carrea et al., 2019; Cole et al., 2019; Gonzalez-Wevar et al., 2019; Rexer-Huber et al., 2019; Baird et al. 2021). In some cases, low levels of dispersal may permit postglacial recolonization, but may fail to provide sufficient gene flow to prevent speciation. Such scenarios are exemplified by the terrestrial Ectemnorhinini weevils and the subtidal limpet genus *Nacella*, which exhibit island-level endemism around the entire Southern Ocean, with divergences dating to the Plio-Pleistocene boundary (~2.58 Mya) and the mid-Pleistocene (0.25-0.6 Mya), respectively (Gonzalez-Wevar et al., 2017; Gonzalez-Wevar et al., 2019; Baird et al., 2021).

At the opposite end of the scale, the highly dispersive southern bull kelp (*Durvillaea antarctica;* a keystone intertidal species), and its associated rafting invertebrate communities, exhibit a ‘northern richness; southern purity’ model whereby disjunct high-latitude sub-Antarctic islands share anomalously low-diversity, circumpolar lineages (Fraser et al., 2009; Nikula et al., 2010; Fraser et al., 2010; Fraser et al., 2011; Fraser et al., 2012; Fraser et al., 2018; Gonzalez-Wevar et al., 2018; Cole et al., 2019). Fraser et al. (2009) hypothesized that these patterns reflect recent passive post-Last Glacial Maximum (LGM) recolonization from ice-free refugia facilitated by circumpolar ocean currents. The intertidal air-breathing limpets *Siphonaria lateralis* and *S. fuegiensis*, which are closely associated with bull kelp, appear to show a more persistent occupation of high-latitude sub-Antarctic islands, as evidenced by their negligible phylogeographic structure suggesting continuing gene flow throughout their range (Gonzalez-Wevar et al., 2018). Intriguingly, the highly mobile southern elephant seal (*Mirounga leonina*), which exhibits a multiregional refugial pattern (de Bruyn et al., 2009) (see also Rexer-Huber et al., 2019), rapidly expanded their range to newly ice-free parts of the Antarctic coast (2500 km from the nearest breeding colony) 8000 years ago before abandoning this new colony 1000 years ago when a grounded ice-sheet returned. These apparently climate-driven patterns are mirrored to some extent by multispecies demographic analyses of high-latitude penguins which reveal concordant evidence for simultaneous and rapid post-LGM population expansions associated with exposure of snow-free ground on which to breed (Cole et al., 2019).

These contrasting biogeographic responses to the Pleistocene glacial-interglacial cycles are underpinned by biology. The gilled *Nacella* limpets can survive subtidally, including on the Antarctic Peninsula (Gonzalez-Wevar et al., 2017; Gonzalez-Wevar et al., 2019), and are thus not as affected by continual ice as more intertidal species like southern bull kelp (despite its ability to passively drift onto Antarctic shores; Fraser et al., 2018), air-breathing *Siphonaria* limpets (Gonzalez-Wevar et al., 2018) or southern elephant seals (de Bruyn et al. 2009).

Blue-eyed shags (*Leucocarbo* spp.) are an ecologically important and highly speciose group of philopatric seabirds exhibiting a circumpolar Southern Ocean distribution, with 17 currently accepted species and subspecies (Kennedy & Spencer, 2014; Rawlence et al., 2017). Many of these taxa are endemic to single island groups (Fig. 1). Additionally, the small number of lineages that breed on mainland coasts typically exhibit strong phylogeographic structure (Calderon et al., 2014; Kennedy & Spencer, 2014; Rawlence et al., 2015; Rawlence et al., 2017). Preliminary genetic research has suggested that blue-eyed shags comprise two widespread, species-rich clades: a sub-Antarctic clade encompassing South America, Antarctica, and high-latitude sub-Antarctic islands; and the other occurring across the New Zealand region (Fig. 1) (Kennedy & Spencer, 2014). Data from the subfossil record and ancient DNA analysis suggests that at least one recently-recognized mainland taxon (*L. chalconotus*) has suffered substantial loss of genetic and geographic diversity since human arrival in mainland New Zealand (Rawlence et al., 2015; Rawlence et al., 2017).

The broad, species-rich biogeography of *Leucocarbo* presents something of a paradox. Specifically, while the wide Southern Ocean distribution of the genus, incorporating numerous isolated islands, implies strong dispersal capacity, the presence of numerous single-island endemics suggests that such dispersal ability is insufficient to prevent widespread speciation. This situation in blue-eyed shags may be somewhat akin to iconic radiations described from remote archipelagic systems such as Hawaii, the Galapagos, and the Canaries (Mendelson & Shaw, 2005; Shaw & Gillespie, 2016) in which the evolution of island endemics has apparently been underpinned by founder speciation (Waters et al., 2013). Unlike these low latitude archipelagos, however, the Southern Ocean islands have been heavily impacted by recent glacial cycles (Fraser et al. 2009; Fraser et al., 2012), suggesting that much of their biotic diversity may have evolved relatively recently.

Based on the distinctive Southern Ocean biogeography of *Leucocarbo* shags, we hypothesize that glacial-interglacial transitions have presented crucial opportunities for rapid circumpolar range expansion and founder speciation in this group. Here we present the first multigene analysis of *Leucocarbo*, that includes all recognized taxa, using a temporal phylogenetic framework to test for rapid radiation of this unusually speciose Southern Ocean assemblage.

## 2. MATERIALS AND METHODS

### 2.1 Specimens, DNA extraction, PCR and sequencing

Tissue, blood, or feathers were obtained from a number of sources, covering the geographic distribution of *Leucocarbo* shags (Appendix One Table S1.1-1.2). Total genomic DNA was extracted using a phenol/chloroform extraction, a 5% Chelex 100 solution or the Qiagen DNeasy Tissue Kit (Walsh et al., 2013; Kennedy & Spencer, 2014). Negative controls were included with each extraction. For the phylogenetic dataset (using a single location per taxon, except for four taxa where two locations were used and one taxon where three locations were used, see Appendix One Table S1.1) DNA was amplified for five mitochondrial genes (12S, overlapping ATPase 8 and 6, ND2, COI) and five nuclear genes (FIB7, PARK7, IRF2, CRYAA, RAPGEF1) (following Kennedy & Spencer, 2014; Kennedy et al., 2019). The phylogenetic dataset’s primer details are shown in Table S1 of Kennedy et al. (2019). To investigate within taxon diversity control region (CR) sequences were amplified for multiple individuals per location (except for two taxa where only a single individual was able to be used, see Appendix One Table S1.2) (following Rawlence et al., 2014). Negative controls were included with each PCR reaction. PCR products were purified using the Ultra-Sep Gel extraction kit (Omega) and sequenced on an Applied Biosystems 3730 capillary sequencer. Newly generated sequences for the phylogenetic and CR datasets were added to, and aligned with, those previously published (Kennedy & Spencer, 2014; Rawlence et al., 2014; Rawlence et al., 2017) (see Appendix One Tables S1.1-S1.2).

### 2.2 Phylogenetic and demographic analysis

The phylogenetic dataset was divided into nine partitions, five nuclear loci and four mitochondrial loci (the overlapping ATPase 8 and 6 were treated as a single ATPase partition). Models of nucleotide substitution were selected using the Akaike Information Criterion of Modeltest 3.7 (Posada & Crandall, 1998). The models selected for each gene region were: HKY + I for 12S (2st + I), GTR + I for ATPase (6st + I), TIM + G for ND2 (6st + G), TIM + I for COI (6st + I), HKY + I for FIB7 (2st + I), HKY for PARK7 (2st), TrN for IRF2 (6st), HKY for CRYAA (2st), and HKY + I for RAPGEF1 (2st + I). We used StarBEAST2 (v. 0.15.13) implemented in BEAST 2.6.3 (Bouckaert et al., 2019) to jointly infer the *Leucocarbo* species tree along with mitochondrial and individual nuclear gene trees. We implemented an analytical population size integration model (Bouckaert et al., 2019), unlinked substitution models for all partitions, linked trees for mitochondrial genes, and a birth-death tree prior. Strict molecular clocks were used; one linked clock for nuclear genes and unlinked clocks for mitochondrial genes. The clock rates for mitochondrial genes were modelled as normal priors with mean substitution rate estimated from rates for terminal nodes 33 to 39 (i.e., the clade *Leucocarbo* belongs to) in Figure S2 of Pacheco et al. (2011) (Appendix One Table S1.3). Mutation rates for individual nuclear genes were modelled with 1/X priors. We ran three independent MCMC chains, each run for 50 million steps, sampling every 5,000 steps. Additionally, to estimate per species population sizes, analyses were rerun using the linear with constant root populations model with the same parameters, but increasing the MCMC chain to 100 million steps, sampling every 10,000 steps. We checked for convergence and sufficient sampling of parameters in Tracer v1.7.1 (Rambaut et al., 2018) and combined individual runs after discarding the first 10% of steps as burn-in in LogCombiner. Maximum clade credibility consensus trees were generated in TreeAnnotator using the median node age. DensiTree v2.2.7 (Bouckaert, 2010) was used to simultaneously visualise all trees post burn-in and generate consensus trees scaled by estimated effective population size.

### 2.3 Ancestral range distribution estimation

We estimated the ancestral range of internal nodes of the *Leucocarbo* species tree using the R package BioGeoBears (Matzke, 2013). We implemented three methods of ancestral state estimation with and without the jump dispersal parameter (J) (Matzke, 2014), and the dispersal probability as a function of the distance parameter (X) (van Dam & Matzke, 2016): dispersal-extinction-cladogenesis (DEC) (Ree & Smith, 2008), dispersal vicariance analysis (DIVA) (Ronquist, 1997), and Bayesian analysis of biogeography (BAYAREA) (Landi et al., 2013). We compared models using corrected Akaike Information Criterion (AICc) and the weighted AICc values. Due to the criticism of +J models (Ree & Sanmartin, 2018), especially regarding statistical comparison to non-+J models, we did not statistically compare models with and without the +J parameter. We split the samples into seven biogeographic areas: N) North America, S) South America (including the Falkland/Malvinas Islands), O) South Orkney Islands, South Georgia and the South Sandwich Islands, A) Antarctic Peninsula, K) Kerguelen, Heard, Marion and Crozet islands, M) Macquarie Island, and Z) New Zealand and the New Zealand sub-Antarctic islands. For models including the +X parameter, pairwise distances between geographic areas were estimated using Google Earth and normalized against the shortest pairwise distance. All analyses were non-stratified with equal transition between areas.

### 2.4 Median Joining Network of mitochondrial control region data

PopArt (Leigh & Bryant, 2015) was used to construct a median joining network of the mitochondrial control region (CR) data. Sites with >5% unidentified states were masked in the analysis.

## 3. RESULTS

### 3.1 Recent divergence and demographic expansion at high latitudes

*Leucocarbo* mitochondrial and nuclear sequences were obtained from across the geographical distribution of blue-eyed shags (and two outgroups, see Appendix One Table S1.1). Of the 8200 characters in the partitioned mitochondrial and nuclear dataset, 7727 were constant, while 282 out of 437 variable characters were parsimony informative.

*Leucocarbo* shags form a strongly supported monophyletic clade diverging from *Nannopterum* 13.3 Mya (HPD 19.8-8.3 Mya) during the mid-Miocene (Fig. 1). All major clades within the species tree show moderate to strong support (0.85-1.0 PP). The rock shag (*L. magellanicus*) is sister to the remainder of the genus, the split occurring 7.5 Mya (HPD 10.9-4.9 Mya) during the late-Miocene, whereas the lineage leading to the Guanay shag (*L. bougainvillii*) diverged much more recently 2.6 Mya (HPD 3.9-1.6 Mya) coincident with the onset of the Pleistocene Ice Age 2.58 Mya (Fig. 1). Within the rock shag there is phylogeographic structure dividing populations from the Atlantic (Argentina) and Pacific (Chile) coasts, dating to 1.3 Mya (HPD 2.2-0.01 Mya) (Fig. 1) in agreement with previously reported mitochondrial data (Calderon et al., 2014) and plumage differences (Rasmussen, 1987). However, microsatellite data suggests recent secondary contact and geneflow from the Pacific to Atlantic coasts (Calderon et al., 2014).

The remaining *Leucocarbo* lineage (leading to all the other extant species) split into two major clades 2.3 Mya (HPD 3.3-1.5 Mya) during the early Pleistocene, shortly after its separation from the Guanay shag lineage (hence the small inter-nodal distance and low posterior probability, PP). The first of these clades contains species from the New Zealand region, while the second comprises taxa from the sub-Antarctic, including South America (and Falkland/Malvinas Islands), Antarctic Peninsula and high latitude sub-Antarctic islands (Fig. 1). Within the sub-Antarctic clade, there are three subclades, the first consisting of species from South America, Falkland/Malvinas Islands, South Georgia and South Orkney Islands; and second (and third) clade comprising taxa from the Antarctica Peninsula, Marion Island, Crozet Island, and Kerguelen, Heard and Macquarie islands, respectively. The Crozet shag (*L. melanogenis*), which inhabits both Crozet and Marion islands, is paraphyletic (Fig. 1).

DensiTree analysis, which shows congruence or conflict among individual gene-tree topologies, shows broad congruence for the major splits in the species tree (Appendix One Fig. S1.1-1.2). However, there are conflicts within the two major clades within *Leucocarbo*, and the relative placement of some taxa (e.g. Bounty Island shag, *L. ranfurlyi* and New Zealand king shag, *L. carunculatus*).

While PP support within the major New Zealand and sub-Antarctic clades is low (*cf*. mitochondrial species tree; Appendix One Fig. S1.1), phylogenetic analyses revealed substantially deeper divergences among temperate *Leucocarbo* lineages relative to those among sub-Antarctic lineages of this genus (Fig. 1). The branch lengths in the New Zealand clade are up to four times longer (up to 2 Mya in the early Pleistocene versus up to 0.5 Mya in the mid-Pleistocene). For instance, while the vast majority of divergence time HPD’s among New Zealand taxa substantially predate the LGM, several sub-Antarctic taxa yielded divergence HPDs that are potentially consistent with post-LGM recolonization (Appendix One Table S1.4).

Demographic analysis based on the species tree shows stable or increasing population size for blue-eyed shags in South America, compared to founder event induced bottlenecks and subsequent population expansions associated with island endemics (Appendix One Fig. S1.3). The finding of relatively short, potentially post-LGM branch lengths in the sub-Antarctic clade (Fig. 1), is also mirrored to some extent by these demographic reconstructions that reveal recent population expansions in numerous sub-Antarctic lineages following major late-Quaternary bottlenecks (Appendix One Fig. S1.3).

In the mitochondrial CR dataset, there were 1026 characters, of which 925 were constant, while 83 out of 101 variable characters were parsimony informative. The median joining network of fast-evolving mitochondrial CR sequences from multiple individuals for the majority of each *Leucocarbo* species (Appendix One Fig. S1.4) shows deep, clear separation of rock and Guanay shags and those from the New Zealand region in contrast to the blue-eyed shags from the sub-Antarctic clade. All species of blue-eyed shags form their own genetic clusters for the CR data, with the exception of the Crozet shag (*L. melanogenis*). There are shallow to deep (e.g. rock shag) levels of genetic variation within blue-eyed shags, with the exception of the Campbell (*L. campbelli*), Auckland (*L. colensoi*), Bounty (*L. ranfurlyi*), and Macquarie (*L. purpurascens*) Island shags, where single haplotypes are present. The blue-eyed shag CR substitution rate of 3.2 s/s/Ma (95% HPD 0.70-6.57; Rawlence et al., 2015) suggests divergence dates broadly consistent with those determined in our phylogenetic analysis of nuclear and mitochondrial DNA markers.

### 3.2 Founder event speciation

The best-supported BioGeoBears evolutionary scenario for blue-eyed shags, based on our phylogenetic analysis, was a jump dispersal (founder-event speciation) – extinction – cladogenesis (DEC+J) model (Fig. 1, Appendix One Table S1.5). This founder-speciation model reconstructed an ancestral geographical area for *Leucocarbo* of South America, and for all major nodes in the phylogeny. The inferred most-recent common ancestor of the two major clades within *Leucocarbo* is from South America, with subsequent circumpolar expansion supported by the Antarctic Circumpolar Current (ACC). The best-supported scenario without a jump parameter was DEC (expansion and speciation) (Appendix One Fig. S1.5, Table S1.5). In contrast to DEC+J, ancestors at deeper nodes in the phylogeny are inferred to have been geographically widespread, with temporally offset *in situ* speciation on different islands.

## 4. DISCUSSION

Genetic analysis of *Leucocarbo* shags reveals a remarkably shallow evolutionary history among endemic lineages breeding at island groups scattered across the vast Southern Ocean, consistent with a recent high-latitude circumpolar radiation. This rapid sub-Antarctic archipelagic expansion contrasts with significantly deeper lineages detected at temperate latitudes in South America and New Zealand that may have acted as glacial refugia. Recent high-latitude expansions are further supported by ancestral demographic and biogeographic reconstructions.

### 4.1 Postglacial circumpolar expansion

The shallow phylogenetic history of sub-Antarctic *Leucocarbo* (Fig. 1) relative to mid-latitude lineages in the New Zealand region (up to 2 Mya) and South America (up to 7.5 Mya) almost certainly reflects the impact of repeated glacial-interglacial cycles during the Pleistocene (2.58 – 0.01 Mya) in underpinning Southern Ocean extinction-recolonization events (Fraser et al., 2009; Nikula et al., 2010; Fraser et al., 2012). Specifically, our analysis detected a four-fold difference in branch lengths between the two major clades of *Leucocarbo* (Fig. 1), with deep phylogenetic histories detected among temperate lineages in putative glacial refugia (e.g., New Zealand islands). This recent expansion history of sub-Antarctic taxa to previously ice-bound, potentially glaciated islands is further highlighted by latitudinal contrasts in demographic reconstructions (Appendix One Fig. S1.3) and echoes recent demographic comparisons of other iconic Southern Ocean seabirds with circumpolar distributions (Trucchi et al., 2014; Cole et al., 2019).

Circumpolar dispersal events (Fig. 1) were likely facilitated by the ACC and westerly winds encircling Antarctica (Fraser et al., 2012). While dating of the shallowest phylogenetic nodes is likely confounded by population-level time-dependant processes (Burridge et al., 2009) (making disentangling the effects of repeated Pleistocene glacial-interglacial cycles difficult), the short internodal distances between far-flung lineages of the Southern Ocean (e.g. Macquarie versus Kerguelen; Antarctic Peninsula versus Marion Island), separated in some instances by ~10,000 km, could correspond to colonization events in several of the recent post-glacial periods. The shortest internodal distances likely reflect post-LGM dispersal (Fig. 1). Indeed, as a proxy for earlier glacial periods, these southern coasts are hypothesized to have been heavily affected by LGM winter sea ice (Ehlers et al., 2011), and in some cases ice and snow cover (Hodgson et al., 2014), which extended north of the current route of the Antarctic Polar Front (Fig. 1; Fraser et al., 2009). In addition to its impact on coastal benthic and intertidal species (Fraser et al., 2009; Nikula et al., 2010; Gonzalez-Wevar et al., 2018), extensive sea ice during glacial maxima is thought to have extirpated Southern Ocean avian lineages that depend on coastal ecosystems (Trucchi et al., 2014; Cole et al., 2019). *Leucocarbo* shags are inshore foragers largely dependent on benthic species which they hunt through pursuit-diving (Cook et al., 2013). Limited by dive depth and foraging range, blue-eyed shags are particularly prone to ecosystem disturbance (Cook et al., 2013; Rawlence et al., 2016), though less so than intertidal species in this context (de Bruyn et al., 2009; Fraser et al., 2009; Gonzalez-Wevar et al., 2018) (as their presence on the seasonally ice-impacted Antarctic Peninsula indicates). Our phylogeographic analyses (Fig. 1) strongly suggest that the New Zealand region and unglaciated South America (including the Falkland/Malvinas Islands) both acted as key glacial refugia, consistent with previous analyses (Austin et al., 2013; Cole et al., 2019; Rexer-Huber et al., 2019).

The inferred postglacial expansion out of South America involved two large-scale, rapid dispersal events: the first from the South American mainland (including Falkland/Malvinas Islands) to the archipelagos of South Georgia and South Orkney Islands (resolving any ambiguity about the origin of the South Orkney Islands shags; Schrimpf et al., 2018); and the other from the mainland to the Antarctic Peninsula and into the high-latitude sub-Antarctic islands (Fig. 1). Founder-takes-all (or density-dependent) processes (Waters et al., 2013) may help to explain some of these idiosyncratic expansion patterns, notably the failure of the South Georgia shag (*L. georgianus*) to colonize the Antarctic Peninsula (where the Antarctic shag *L. bransfieldensis* breeds), and the absence of close relatives of the Macquarie Island shag (*L. purpurascens*) in the New Zealand region, a mere 500km downwind. However, more complicated dispersal scenarios cannot be discounted given the conflict among gene trees (due to incomplete lineage sorting and/or gene-flow), and between the species tree (Fig. 1) and the mitochondrial CR data (Appendix One Fig. S1.4). Indeed, the South American and Falkland/Malvinas Island forms of the imperial shag (*L. atriceps*, sometimes treated as subspecies *atriceps* and *albiventer*, respectively, e.g. Nelson, 2005) are paraphyletic (in comparison to the monophyly exhibited by the other *Leucocarbo* spp., see Appendix One Fig. S1.4), and no doubt represent the same taxon (see also Calderon et al., 2014). This situation makes it difficult to infer whether blue-eyed shags that dispersed to other islands in the Southern Ocean and the Antarctic continent came from the South American mainland or the nearby Falkland/Malvinas Islands. Regardless, like *Leucocarbo* species in the New Zealand clade, this postglacial dispersal is characterized by the genetic signatures of founder-events and associated population bottlenecks followed by subsequent population expansions.

### 4.2 Rapid founder speciation

Evolutionary biogeographic reconstructions strongly support a founder-speciation model (jump dispersal) for all major nodes of the *Leucocarbo* phylogeny. Indeed, the ‘jump’ parameter is important in modelling founder-event speciation events that often characterize remote archipelagic species (Matzke, 2014; see also Ree & Sanmartin, 2018). By contrast, the best-supported alternative evolutionary scenario lacking such a jump parameter requires widespread ancestors at deeper nodes, which seems biogeographically implausible in this region (Appendix One Figure S1.5, Table S1.4).

Founder speciation represents a fascinating feature of archipelagic biogeography globally (Mendelson & Shaw, 2005; Shaw & Gillespie, 2016), and such phenomena may be heavily influenced by density-dependent processes (Waters et al., 2013; Shaw & Gillespie, 2016). While many of the best-known archipelagic radiations have been detected in tropical systems that have been relatively sheltered from global climatic shifts, our study suggests that such founder-speciation processes may be important even in the vast Southern Ocean (Trucchi et al., 2014; Cole et al., 2019; Baird et al., 2021), a system heavily impacted by geologically recent glacial cycles. Indeed, the apparent pace of colonization and speciation, including previously unrecognized founder events, in Southern Ocean *Leucocarbo* may approach that of the most rapid radiations elsewhere in the globe (Kornfield, 1978; Mendelson & Shaw, 2005; Momigliano et al., 2017; Puntambekar et al., 2020; Turbek et al., 2021). We note that a recent field guide to seabirds (Harrison et al., 2021) depicts significant differences in adult breeding plumage and skin colouration (e.g. extent of white feathering on the cheek, gular colour, size and colour of the carunculation above the bill) among the various species. These differences, composites of details observable in recent detailed digital photographs (Peter Harrison pers. comm.), are likely to be critical in specific-mate recognition.

The Crozet Island shag (*L. melanogenis*), originally described from Crozet Island, but also seemingly present on Marion Island (Blyth, 1860; Moseley, 1879; Alexander, 1928; Holgersen, 1945; Crawford, 1952; Rand, 1954; Rand, 1956; Crawford et al., 2003), is paraphyletic in our analyses (Fig. 1, Appendix One Fig. S1.4), and is possibly the consequence of an unrecognized founder event on Marion Island. Considering the observed plumage differences between blue-eyed shags from Marion and Crozet Islands (Rand, 1954), and phylogeographic structure within *Leucocarbo*, we suggest that the Marion Island population may represent a new undescribed taxon.

It is not clear, however, how *Leucocarbo* shags managed to colonize remote island groups, separated by thousands of kilometres, across an expanse of empty ocean. Shag biology suggests a relatively low dispersal capacity. Like other species in Phalacrocoracidae, *Leucocarbo* shags have a partially wettable plumage that decreases the layer of insulating air and reduces the energetic costs of diving (Gremillet et al., 2005; Cook & Leblanc, 2007). Blue-eyed shags would not be expected to withstand days or weeks at sea in cold Southern Ocean waters, although facultative hypothermia observed in these shags might enable some tolerance to near-freezing sea-water temperatures (Bevan et al., 1997). Flight performance in this group is also rather limited compared to other avian lineages (Pennycuick, 1989), in part due to their dense bones, which aid in diving. A several thousand-kilometre non-stop flight across the Southern Ocean would therefore seem unlikely, unless driven by some particularly strong and sustained westerly wind and current system. Furthermore, due to the depth of the Southern Ocean, access to benthic prey would be impossible during stopovers on the water surface, although *Leucocarbo* shags are flexible enough to occasionally forage on pelagic fish species (Cook et al., 2013).

It is possible that blue-eyed shags dispersed eastwards from South America following the frontline of the receding sea-ice after glacial maxima and were thus able to approach recently ice-freed remote sub-Antarctic islands more easily. Regardless of the colonization or recolonization pathway, it is a distinct possibility that the sub-Antarctic islands were colonized by small groups of shags (10-20 birds), rather than by lone individuals, as blue-eyed shags will readily forage or travel in groups (Cook et al., 2013).

### 4.3 Sentinels of change

Intriguingly, *Leucocarbo* shags exhibit a mixture of biological characteristics seen in both poorly and highly dispersive taxa. The phylogeographic pattern of multiple deep lineages in the New Zealand region and South America, versus shallow lineages in Antarctica and high-latitude sub-Antarctic islands, provides a clear signature of rapid expansions from ice-free refugia into previously glaciated and/or ice-bound areas (Fraser et al., 2012), and strongly indicates a sensitivity to climate change. The strong philopatric behaviour of *Leucocarbo* shags has resulted in high levels of endemism, a pattern that might be expected for less dispersive taxa (Gonzalez-Wevar et al., 2017; Gonzalez-Wevar et al., 2019; Rexer-Huber et al., 2019; Baird et al., 2021). The circumpolar distribution of blue-eyed shags, and their highly dynamic evolutionary history in relation to climate change (Calderon et al., 2014; this study) and anthropogenic impacts (Rawlence et al., 2014; Rawlence et al., 2015; Rawlence et al., 2017), potentially make *Leucocarbo* a strong sentinel of past and ongoing Southern Ocean ecosystem change.

## Supporting information

Rawlence et al. Suppl. Info.

## ACKNOWLEDGEMENTS

This work was supported with funding from the University of Otago. We gratefully acknowledge the following people and institutions for kindly providing samples for the DNA sequences new in this study: J. Amey, C. Fraser, G. Frisk, C. González-Wevar, J. Fyfe, D. Lijtmaer, K. Marchant, I. Nishiumi, G. Nunn, D. Onley, R. Palmer, E. Poulin, R. Schuckard, A. Scolaro, P. Sweet, T, Webster; the American Museum of Natural History, Australian National Wildlife Collection (CSIRO), Otago Museum, Museo Argentino de Ciencias Naturales, Museum of New Zealand Te Papa Tongarewa, New Zealand Department of Conservation, and the Swedish Museum of Natural History. We thank the IPEV (Insitut Polaire Francais) for support to Program N° 394. We would also like to recognize all those people and institutions that kindly provided samples for the previously published sequences also used in this study. We thank Peter Harrison for discussions about the details of the plumage and colouration of breeding blue-eyed shags.

## DATA ACCESSIBILITY

Data and scripts to reproduce the results of this study are available on Data Dryad: XXXXX. Sequences have been uploaded to GenBank (see Appendix One Tables S1.1-S1.2).

## BIOSKETCH

**Nic Rawlence** is a biologist who uses genetic tools to study evolutionary processes. His work is primarily focused on reconstructing the impacts of climate change and humans on ecosystems through time.

## Author Contributions

NJR, JMW, HGS, CET and MK developed the study concept. NJR, ATS, JMW, HGS and MK designed the project and wrote the initial manuscript. TRC, C-AB, LJN, PGR, LC, PQ, JFM, NR and RAP collected and/or provided samples. CET, TMK and MK generated the sequence data. ATS, LS, NJR, LD and MK generated results. TRC, LJN, PGR, LC and RAP helped design sampling and project directions. All authors contributed to the final manuscript.

