## Supplementary material for "Rapid radiation of Southern Ocean shags in response to receding sea ice": Rawlence et al. Suppl. Info.

**Supplementary Material: Appendix One**

Nicolas J. Rawlence^1^, Alexander T. Salis^1, 2^, Hamish G. Spencer^1^, Jonathan M. Waters^1^, Lachie Scarsbrook^1^, Richard A. Phillips^3^, Luciano Calderón^4^, Timothée R. Cook^5^, Charles-André Bost^6^, Ludovic Dutoit^1^, Tania M. King^1^, Juan F. Masello^7^, Lisa J. Nupen^8^, Petra Quillfeldt^7^, Norman Ratcliffe^3^, Peter G. Ryan^5^, Charlotte E. Till^1, 9^, Martyn Kennedy^1^

^1^ Department of Zoology, University of Otago, Dunedin, New Zealand.

^2^ Australian Centre for Ancient DNA, University of Adelaide, South Australia, Australia.

^3^ British Antarctic Survey, Natural Environment Research Council, United Kingdom.

^4^ Instituto de Biología Agrícola de Mendoza (IBAM, CONICET-UNCuyo), Argentina.

^5^ FitzPatrick Institute of African Ornithology, Department of Biological Sciences, University of Cape Town, South Africa.

^6^ CEBC-CNRS, UMR 7372, 405 Route de Prissé la Charrière, 79360 Villiers en Bois, France.

^7^ Justus Liebig University, Giessen, Germany.

^8^ Organisation for Tropical Studies, Skukuza, South Africa.

^9^ School of Human Evolution and Social Change, Arizona State University, Arizona, USA.

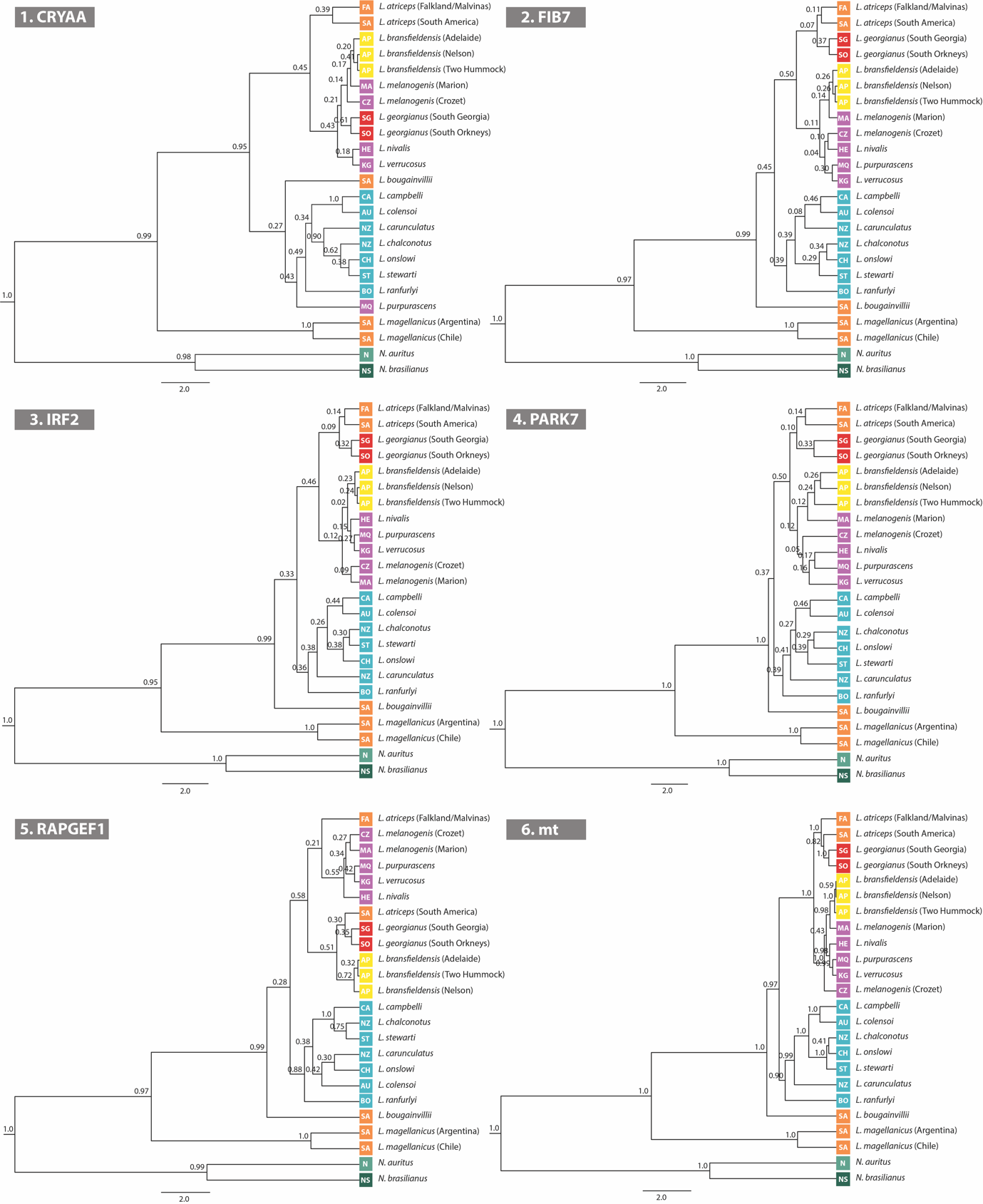

**Figure S1.1** Individual nuclear and mitochondrial gene trees of *Leucocarbo* shags. Node values are Bayesian posterior probability support. Colours and abbreviations are as follows: orange: South America; purple: high-latitude sub-Antarctic islands; yellow: Antarctic Peninsula; blue: New Zealand region; olive: North America; dark green: South America; FA: Falkland/Malvinas Islands; SA: South America; SG: South Georgia; SO: South Orkney Islands: AP: Antarctic Peninsula; MA: Marion Island; CZ: Crozet Island; HE: Heard Island; MQ: Macquarie Island; KG: Kerguelen Islands; CA: Campbell Island; AU: Auckland Island; NZ: mainland New Zealand; ST: Stewart Island; CH: Chatham Islands; BO: Bounty Islands; N: North America; NS: North and South America.

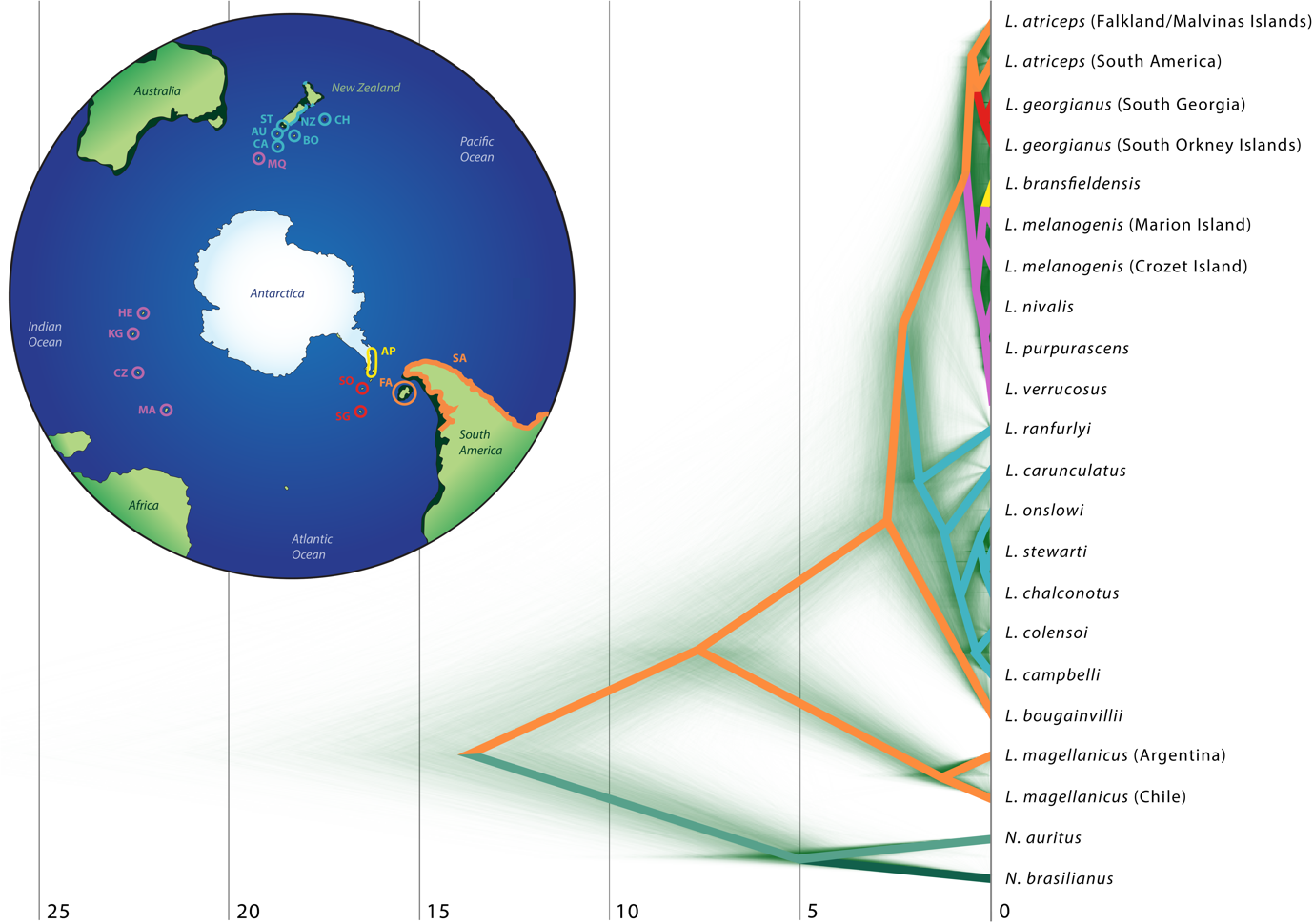

**Figure S1.2** DensiTree analysis of the *Leucocarbo* SpeciesTree showing congruence and conflict between nuclear and mitochondrial gene trees (opaque green). Vertical bars represent divergence times in millions of years. Branch colours are as follows: orange: South America; purple: high-latitude sub-Antarctic islands; yellow: Antarctic Peninsula; blue: New Zealand region; olive: North America; dark green: South America. Divergence times are in millions of years.

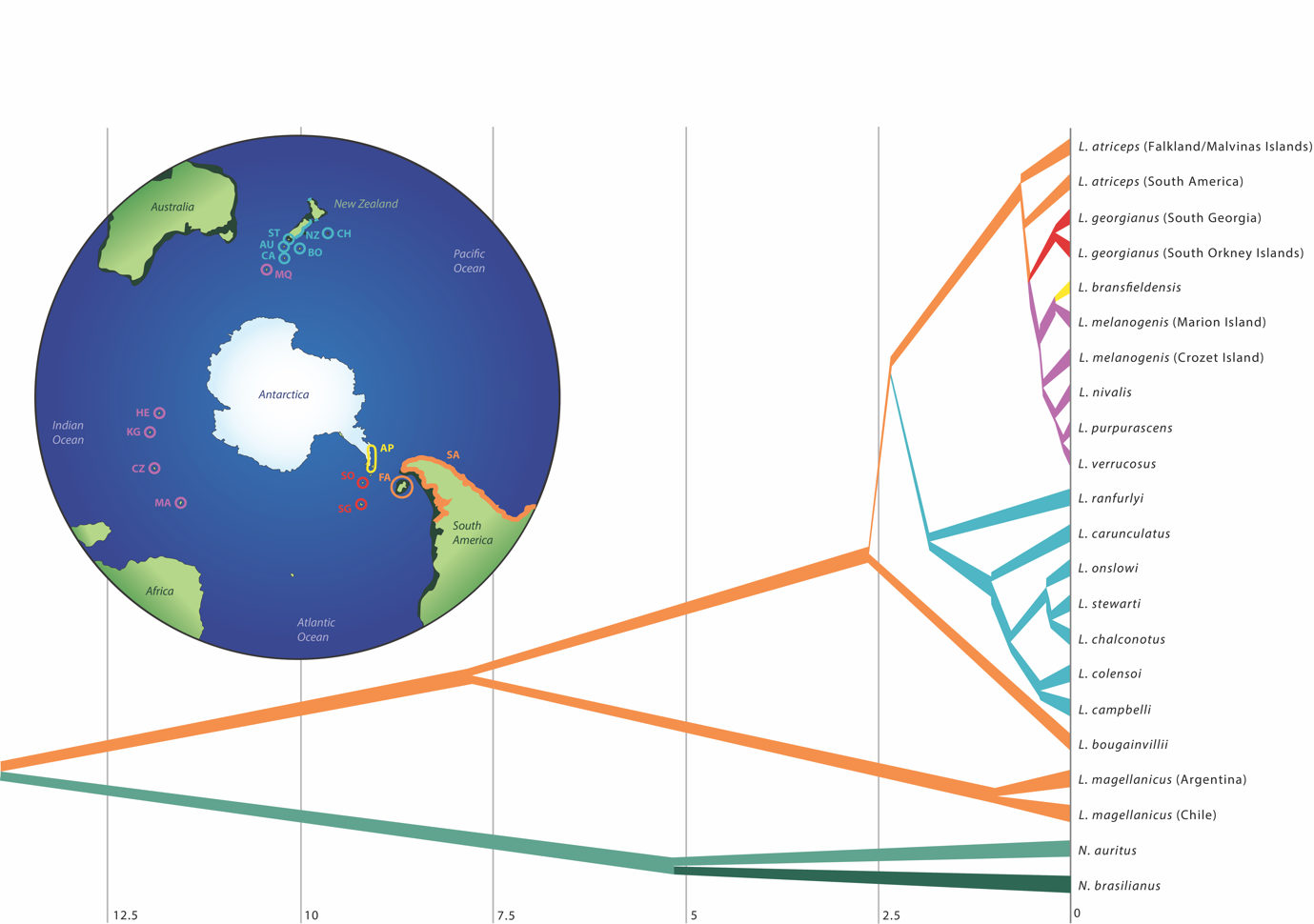

**Figure S1.3** Demographic analysis of *Leucocarbo* shags based on the reconstructed species tree. Branch widths represent relative population size estimates (e.g., narrow branches represent bottlenecks or small founder populations, while widening branches represent population expansions) based on the generation time of blue-eyed shags. The map depicts the geographic distribution of blue-eyed shags. Colours and abbreviations are as follows: orange: South America; purple: high-latitude sub-Antarctic islands; yellow: Antarctic Peninsula; blue: New Zealand region; olive: North America; dark green: South America; FA: Falkland/Malvinas Islands; SA: South America; SG: South Georgia; SO: South Orkney Islands: AP: Antarctic Peninsula; MA: Marion Island; CZ: Crozet Island; HE: Heard Island; MQ: Macquarie Island; KG: Kerguelen Islands; CA: Campbell Island; AU: Auckland Island; NZ: mainland New Zealand; ST: Stewart Island; CH: Chatham Islands; BO: Bounty Islands; N: North America; NS: North and South America.

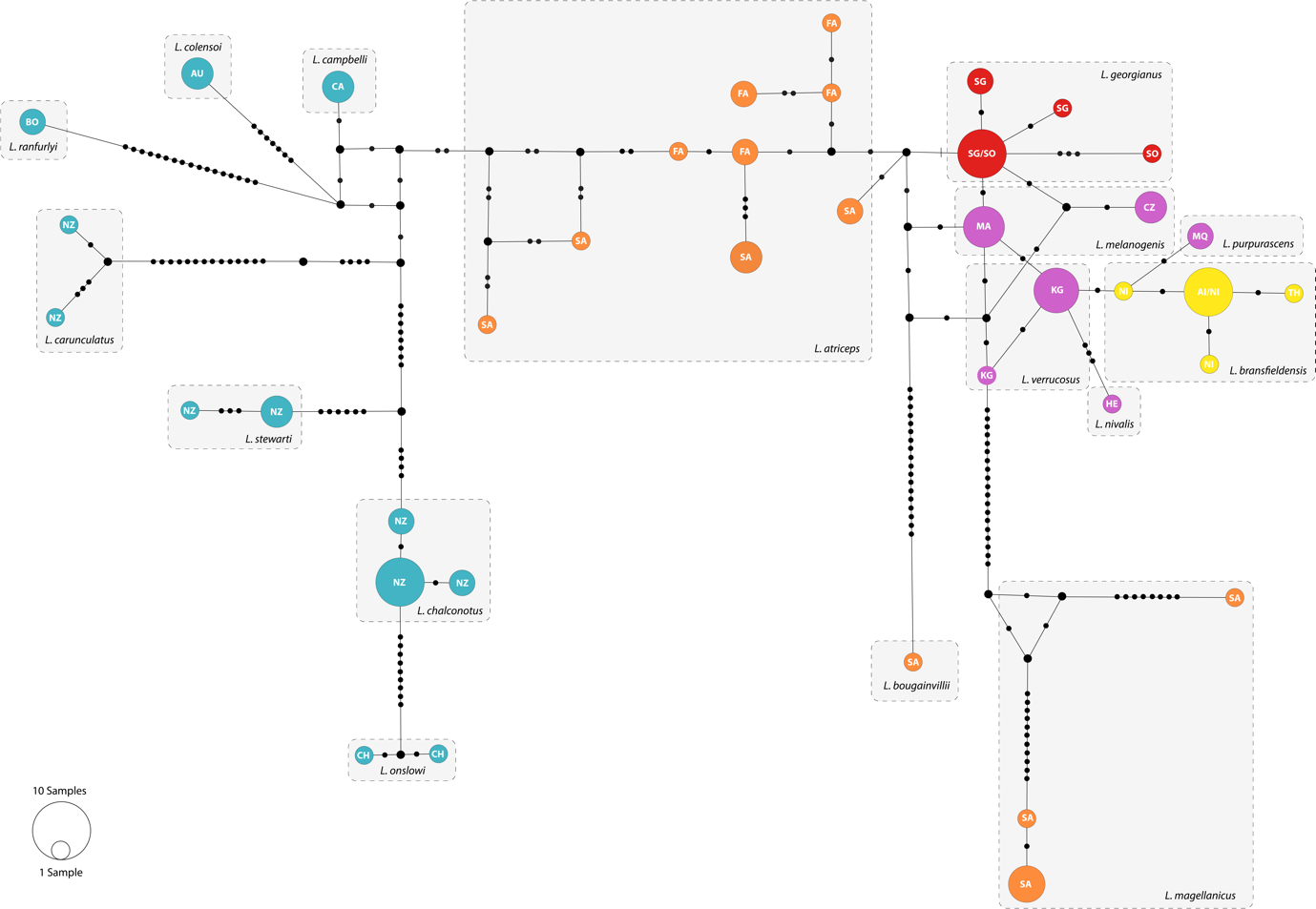

**Figure S1.4** Median joining haplotype network of Control Region from *Leucocarbo* shags. Circles represent unique haplotypes, with the size of the circle proportional to the number of individuals sharing that haplotype. Black circles represent inferred intermediate haplotypes. Colours and abbreviations are as follows: orange: South America; purple: high-latitude sub-Antarctic islands; yellow: Antarctic Peninsula; blue: New Zealand region; FA: Falkland/Malvinas Islands; SA: South America; SG: South Georgia; SO: South Orkney Islands: AP: Antarctic Peninsula; MA: Marion Island; CZ: Crozet Island; HE: Heard Island; MQ: Macquarie Island; KG: Kerguelen Islands; CA: Campbell Island; AU: Auckland Island; NZ: mainland New Zealand; ST: Stewart Island; CH: Chatham Islands; BO: Bounty Islands.

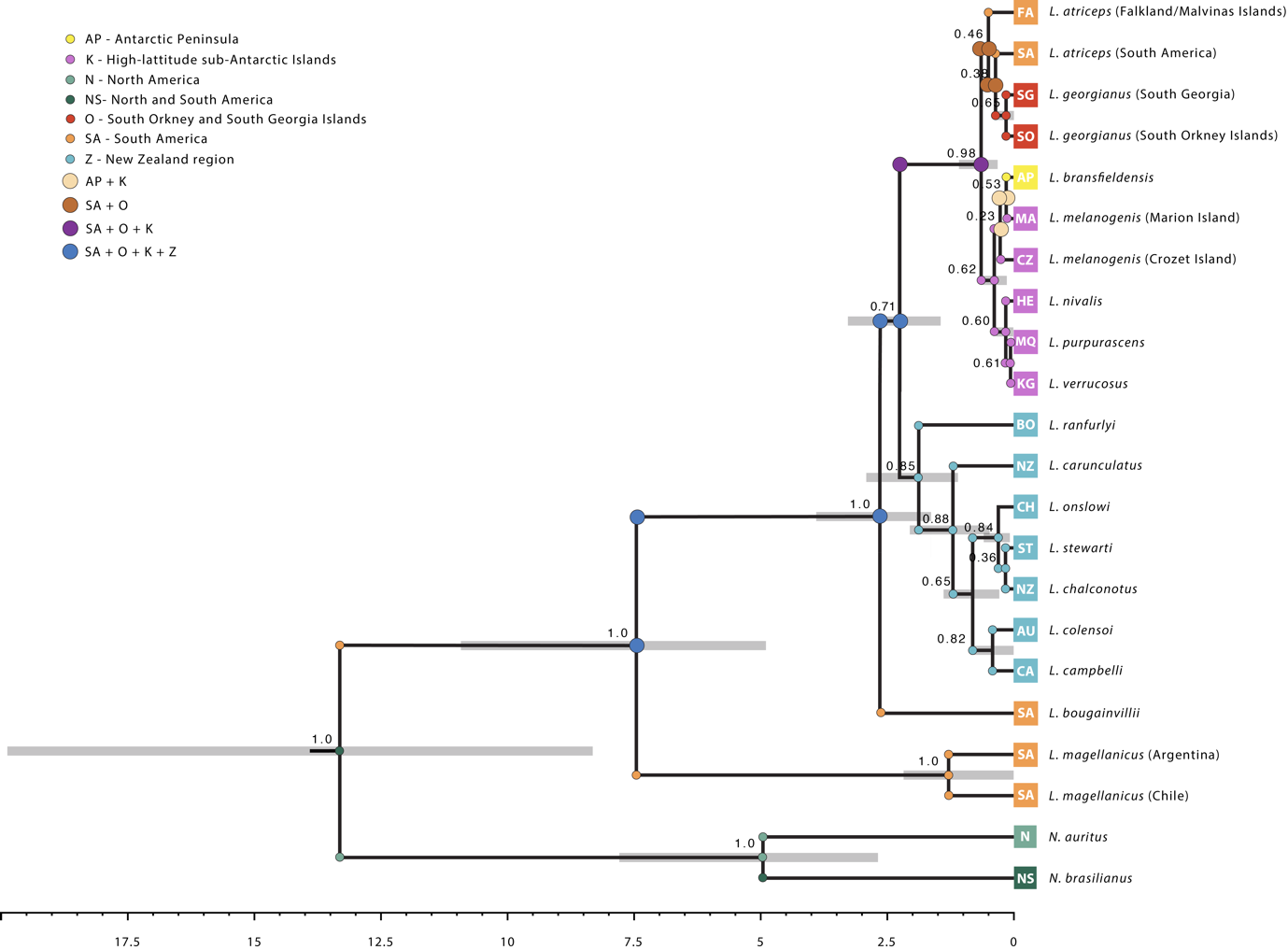

**Figure S1.5** Evolutionary history of blue-eyed shags (*Leucocarbo* spp.) The mitochondrial and nuclear DNA species tree, based on 8.2 kilobases of DNA sequence data (five mitochondrial and five nuclear genes) depicts the evolution of blue-eyed shags. Node bars on the phylogeny are 95% HPD of divergence times. Node values are Bayesian posterior probability support. Node circles are the ancestral state reconstruction of the geographic distribution based on the DEC model. Colours and abbreviations are as follows: orange: South America; purple: high-latitude sub-Antarctic islands; yellow: Antarctic Peninsula; blue: New Zealand region; olive: North America; dark green: South America; FA: Falkland/Malvinas Islands; SA: South America; SG: South Georgia; SO: South Orkney Islands: AP: Antarctic Peninsula; MA: Marion Island; CZ: Crozet Island; HE: Heard Island; MQ: Macquarie Island; KG: Kerguelen Islands; CA: Campbell Island; AU: Auckland Island; NZ: mainland New Zealand; ST: Stewart Island; CH: Chatham Islands; BO: Bounty Islands; N: North America; NS: North and South America.

**Table S1.1** GenBank accession numbers for each maker in the *Leucocarbo* phylogenetic dataset.

| Taxon | 12S | ATPase-8 & -6 | ND2 | COI | FIB7 | PARK7 | IRF2 | CRYAA | RAPGEF1 |
| --- | --- | --- | --- | --- | --- | --- | --- | --- | --- |
| *Nannopterum auritus* | AY009328 | AY009352 | KM066483 | EF101683 | KM066279 | KM066318 | KM066359 | KM066399 | KM066436 |
| *Nannopterum brasilianus* | AY009336 | AY009360 | KM066484 | KM066519 | KM066280 | KM066319 | KM066360 | KM066400 | KM066437 |
| *Leucocarbo atriceps* (Falkland/Malvinas Islands) | KM066252 | KM066452 | KM066466 | KM066502 | KM066264 | KM066300 | KM066341 | KM066383 | KM066420 |
| *Leucocarbo atriceps* (South America) | AY009342 | AY009366 | KM066467 | KM066503 | KM066265 | KM066301 | KM066342 | KM066384 | KM066421 |
| *Leucocarbo bougainvillii* | AY009330 | AY009354 | KM066468 | KM066504 | **—** | KM066302 | KM066343 | **—** | **—** |
| *Leucocarbo bransfieldensis* (Adelaide Island) | **AB######** | **AB######** | **AB######** | **AB######** | **AB######** | **—** | **AB######** | **AB######** | **AB######** |
| *Leucocarbo bransfieldensis* (Nelson Island) | KM066253 | KM066453 | KM066469 | KM066505 | KM066266 | KM066303 | KM066344 | KM066385 | KM066422 |
| *Leucocarbo bransfieldensis* (Two Hummock Island) | GU445904 | KM066454 | KM066470 | KM066506 | KM066267 | KM066304 | KM066345 | KM066386 | KM066423 |
| *Leucocarbo campbelli* | AY009325 | AY009349 | KM066471 | KM066507 | KM066268 | KM066305 | KM066346 | KM066387 | KM066424 |
| *Leucocarbo carunculatus* | KM066254 | KM066455 | KM066472 | KM066508 | KM066269 | KM066306 | KM066347 | KM066388 | KM066425 |
| *Leucocarbo chalconotus* | **AB######** | **AB######** | **AB######** | **AB######** | **AB######** | **AB######** | **AB######** | **AB######** | **AB######** |
| *Leucocarbo colensoi* | GU445901 | GU445907 | KM066474 | KM066510 | KM066271 | KM066308 | KM066349 | KM066390 | KM066427 |
| *Leucocarbo georgianus* (Bird Island) | GU445903 | GU445909 | KM066475 | KM066511 | KM066272 | KM066309 | KM066350 | KM066391 | KM066428 |
| *Leucocarbo georgianus* (Signy Island) | **AB######** | **AB######** | **AB######** | **AB######** | **AB######** | **AB######** | **AB######** | **AB######** | **AB######** |
| *Leucocarbo magellanicus* (Argentina) | **AB######** | **AB######** | **AB######** | **AB######** | **AB######** | **AB######** | **AB######** | **AB######** | **AB######** |
| *Leucocarbo magellanicus* (Chile) | AY009335 | AY009359 | KM066476 | KM066512 | KM066273 | KM066310 | KM066351 | KM066392 | KM066429 |
| *Leucocarbo melanogenis* (Crozet Island) | **AB######** | **AB######** | **AB######** | **AB######** | **AB######** | **AB######** | **AB######** | **AB######** | **AB######** |
| *Leucocarbo melanogenis* (Marion Island) | KM066255 | KM066456 | KM066477 | KM066513 | KM066274 | KM066311 | KM066352 | KM066393 | KM066430 |
| *Leucocarbo nivalis* | KM066256 | KM066457 | KM066478 | KM066514 | **—** | KM066312 | KM066353 | **—** | **—** |
| *Leucocarbo onslowi* | AY009327 | AY009351 | KM066479 | KM066515 | KM066275 | KM066313 | KM066354 | KM066394 | KM066431 |
| *Leucocarbo purpurascens* | AY009334 | AY009358 | KM066480 | KM066516 | KM066276 | KM066314 | KM066355 | KM066395 | KM066432 |
| *Leucocarbo ranfurlyi* | GU445902 | GU445908 | KM066481 | KM066517 | KM066277 | KM066315 | KM066356 | KM066396 | KM066433 |
| *Leucocarbo stewarti* | AY009344 | AY009368 | KM066473 | KM066509 | KM066270 | KM066307 | KM066348 | KM066389 | KM066426 |
| *Leucocarbo verrucosus* | **AB######** | **AB######** | **AB######** | **AB######** | **AB######** | **AB######** | **AB######** | **AB######** | **AB######** |

- Sequence not included for this marker. Two taxa (*bougainvillii* and *magellanicus*, Argentina) are missing sequence for the barcoding region of COI (see Table S1 in Kennedy et al. (2019) for primer information, and Kennedy and Spencer (2014) for the PCR conditions used).

**Table S1.2** Taxa used in the analysis of the Control Region dataset and GenBank accession numbers.

| Taxa | Voucher information/source or holder of sample | | Collection location(s) | Control Region GenBank accession numbers |
| --- | --- | --- | --- | --- |
| Imperial Shag  *Leucocarbo atriceps* (Falkland/Malvinas Islands) | | Unvouchered sample (Falkland), Martyn Kennedy;  Unvouchered samples (CO.07: 124, 200, 225, 226, 237, 240), Petra Quillfeldt/Juan Masello | Rockhopper Point, Sea Lion Island, Falkland Islands;  New Island, Is. Malvinas/Falklands | AB######; AB######; AB######; AB######; AB######; AB######; AB###### |
| *Leucocarbo atriceps* (South America) | | Unvouchered samples (Scol1 and GN5), Martyn Kennedy;  Samples MACN-Or-ct 3379-3382 and 3398 Museo Argentino de Ciencias Naturales (Luciano Calderón) | Puerto Madryn, Argentina and Chile;  Punta León, Provincia de Chubut, Argentina | AB######; AB######; AB######; AB######; AB######; AB######; AB###### |
| Guanay Shag  *Leucocarbo bougainvillii* | | Unvouchered sample (GN7), Martyn Kennedy and sample DOT 3095, American Museum of Natural History | Chile and Región de Tarapacá, Chile | AB###### |
| Antarctic Shag  *Leucocarbo bransfieldensis* (Nelson Island) | | Sample DOT 10287, American Museum of Natural History (sample labeled as *atriceps*, presumed *bransfieldensis* from sampling location); | Captive, originally from Nelson Island, South Shetland Islands, Antarctica; | AB######; AB######; AB######; AB######; AB######; AB###### |
|  | | Samples MACN-Or-ct 5775, 5781 and 5784-5786 Museo Argentino de Ciencias Naturales (Luciano Calderón); | Isla Nelson, Antártida; |  |
| *Leucocarbo bransfieldensis* (Two Hummock Island) | | Sample NRM 896296, Swedish Museum of Natural History (sample labeled as *atriceps*, presumed *bransfieldensis* from sampling location); | Gerlache Strait, Two Hummock Island, Antarctica; | AB###### |
| *Leucocarbo bransfieldensis* (Adelaide Island) | | Unvouchered samples (AI2, AI3, AI5), Richard Phillips | Adelaide Island, Antarctica | AB######; AB######; AB###### |
| Campbell Island Shag  *Leucocarbo campbelli* | | Unvouchered samples (Campbell1 [=CIS1]-Campbell3), Martyn Kennedy | Campbell Island | KJ189964; AB######; AB###### |
| New Zealand King Shag  *Leucocarbo carunculatus* | | Unvouchered samples (King Shag 1), Martyn Kennedy and (NRO303), Nic Rawlence | Duffers Reef and Port Gore, Marlborough Sounds, New Zealand | AB######; AB###### |
| Otago Shag  *Leucocarbo chalconotus* | | Unvouchered samples (SISB1, SISL1, SISL2, SIS11, SISK1, SISO1, SISL4, and SISL5), Martyn Kennedy and (AMNZ: Tax04-85, OM: Av8958 and JFSIS), Nic Rawlence | Oamaru, Wharekakahu Island, Wharekakahu Island, Allen's Beach, Karitane, Warrington Beach, Aramoana, Long Beach Cave, Cooks Head, Sandfly Bay and St Clair, New Zealand | KJ189977; KJ189980; KJ189981; KJ189975; KJ189978; KJ189979; KJ189976; KJ189982; KJ189983; KJ189984; AB###### |
| Auckland Island Shag  *Leucocarbo colensoi* | | Unvouchered samples (AIS1, AISH and AISW), Martyn Kennedy | Auckland Islands; Port Ross, Ewing Island; and North East Cape, Enderby Island | KJ189963; AB######; AB###### |
| South Georgia Shag  *Leucocarbo georgianus* (Bird Island) | | Unvouchered samples (RPh1-6), Richard Phillips | Bird Island, South Georgia | AB######; AB######; AB######; AB######; AB######; AB###### |
| *Leucocarbo georgianus* (Signy Island) | | Unvouchered samples (SI2, SI6, SI14, SI15, SI21), Norman Ratcliffe | Signy Island, South Orkney Islands | AB######; AB######; AB######; AB######; AB###### |
| Rock (or Magellanic) Shag  *Leucocarbo magellanicus* (Argentina) | | Samples MACN-Or-ct 4424-4428 Museo Argentino de Ciencias Naturales (Luciano Calderón) | Punta Loma, Puerto Madryn, Provincia de Chubut, Argentina; | AB######; AB######; AB######; AB######; AB######; AB###### |
| *Leucocarbo magellanicus* (Chile) | | Unvouchered sample (GN6), Martyn Kennedy | Chile |  |
| Crozet Shag (Crozet)  *Leucocarbo melanogenis* | | Unvouchered samples (CS3-5), Timothée Cook | Crozet Island | AB######; AB######; AB###### |
| Crozet Shag (Marion)  *Leucocarbo melanogenis* | | Unvouchered samples (Crozet), Martyn Kennedy and (CSM1-4) Timothée Cook | Marion Island | AB######; AB######; AB######; AB######; AB###### |
| Heard Shag  *Leucocarbo nivalis* | | Sample ANWC:Birds:B49825, Australian National Wildlife Collection, CSIRO | Heard Island | AB###### |
| Chatham Island Shag  *Leucocarbo onslowi* | | Unvouchered samples (CISB1, CISB2), Martyn Kennedy | Chatham Island | KJ189966; KJ189965 |
| Macquarie Shag  *Leucocarbo purpurascens* | | Unvouchered samples (11 and 20), Martyn Kennedy | Macquarie Island | AB######; AB###### |
| Bounty Island Shag  *Leucocarbo ranfurlyi* | | Unvouchered samples (B81 and B83), Martyn Kennedy | Bounty Island | AB######; AB###### |
| Foveaux Shag  *Leucocarbo stewarti* | | Unvouchered samples (SISC1, SISC2, SISE1 and SISL6), Martyn Kennedy | Stewart Island, Southland, Southland and Boulder Beach, New Zealand | AB######; AB######; AB######; AB###### |
| Kerguelen Shag  *Leucocarbo verrucosus* | | Unvouchered samples (KIS1, KIS2, KIS4 and KIS5), Martyn Kennedy and (KS1, KS3 and KS7), Timothée Cook | Kerguelen Island | AB######; AB######; AB######; AB######; AB######; AB######; AB###### |

*Note*: The terms Cormorant and Shag are often used interchangeably for many of these taxa (and depends on local common usage). The unvouchered samples (with the exceptions of the AMNH samples labelled GN5, GN6 and GN7, which were obtained prior to their having accession numbers and, thus, could be any one the samples for their taxon from the AMNH collection) were not sourced from museums and include tissue, blood, and feather samples as well as extracted DNA.

**Table S1.3** Mean rate and standard deviation for the different mitochondrial partitions used in StarBEAST2 analyses in BEAST2. Values were obtained by averaging rate estimates across the terminal nodes (33-39) encompassed by node 23 from Pacheco et al. (2011).

|  | Mean | Standard Deviation |
| --- | --- | --- |
| ND2 | 0.00388 | 0.0013 |
| COX1 | 0.00232 | 0.0007 |
| ATPase | 0.0029 | 0.0011 |
| 12S | 0.00145 | 0.0011 |

**Table S1.4** Divergence times from the *Leucocarbo* SpeciesTree. Colours correspond to those in Figure 1. n/a indicates poorly supported nodes for which 95% HPD intervals were not calculated.

| **Species divergence** | **Mean divergence time** | **95% HPD** |
| --- | --- | --- |
| Split *Leucocarbo* | 13.3 Mya | 19.8-8.3 Mya |
| Split *L. magellanicus* | 7.5 Mya | 10.9-4.9 Mya |
| Split *L. magellanicus* Argentina vs Chile | 1.3 Mya | 2.2 Mya – 10 Kya |
| Split L. *bougainvillii* | 2.6 Mya | 3.9-1.6 Mya |
| Split New Zealand clade | 2.3 Mya | 3.3-1.5 Mya |
| Split *L. ranfurlyi* | 1.9 Mya | 2.9-1.1 Mya |
| Split *L. carunculatus* | 1.2 Mya | 2.1 Mya – 500 Kya |
| Split *L. colensoi* + *campbelli* | 800 Kya | 1.4 Mya – 300 Kya |
| Split *L. colensoi* vs *campbelli* | 400 Kya | 800-0 Kya |
| Split *L. onslowi* | 300 Kya | 300-80 Kya |
| Split *L. stewarti* vs *chalconotus* | 160 Kya | n/a |
| Split Antarctic/sub-Antarctic clade | 600 Kya | 1.1 Mya – 300 Kya |
| Split *L. nivalis* + *purpurascens* + *verrucosus* | 400 Kya | 700-200 Kya |
| Split *L. nivalis* | 200 Kya | 300-10 Kya |
| Split *L. purpurascens* vs *verrucosus* | 70 Kya | 200-0 Kya |
| Split *L. melanogenis* (Crozet Island) | 300 Kya | n/a |
| Split *L. bransfieldensis* vs *melanogenis* (Marion Island) | 200 Kya | 300-0 Kya |
| Split *L. atriceps* (Falkland/Malvinas Islands) | 500 Kya | n/a |
| Split *L. atriceps* (South America) | 400 Kya | n/a |
| Split *L. georgianus* South Georgia vs South Orkney Islands | 200 Kya | 400-0 Kya |

**Table S1.5** Results from BioGeoBEARS analysis, including likelihood (LnL), corrected Aikaike Information Criterion (AICc) and the associated weights (AICc_wt). Models with and without the jump (J) parameter were run separately and not statistically compared.

|  | No jump parameter | | | Jump parameter (+J) | | |
| --- | --- | --- | --- | --- | --- | --- |
|  | LnL | AICc | AICc_wt | LnL | AICc | AICc_wt |
| DEC | -28.76 | 62.15 | 0.49 | -22.48 | 52.29 | 0.64 |
| DEC+x | -27.9 | 63.14 | 0.3 | -22.42 | 55.19 | 0.15 |
| DIVALIKE | -29.81 | 64.26 | 0.17 | -24.35 | 56.03 | 0.098 |
| DIVALIKE+x | -29.76 | 66.85 | 0.046 | -24.16 | 58.67 | 0.026 |
| BAYAREALIKE | -46.52 | 97.67 | 9.40E-09 | -24.67 | 56.67 | 0.071 |
| BAYAREALIKE+x | -46.43 | 100.2 | 2.70E-09 | -24.61 | 59.57 | 0.017 |
